# Mitochondrial Pyruvate Carrier Inhibition Attenuates Hepatic Stellate Cell Activation and Liver Injury in a Mouse Model of Metabolic Dysfunction-associated Steatotic Liver Disease

**DOI:** 10.1101/2023.02.13.528384

**Authors:** Mohammad Habibi, Daniel Ferguson, Sophie J. Eichler, Mandy M. Chan, Andrew LaPoint, Trevor M. Shew, Mai He, Andrew J. Lutkewitte, Joel D. Schilling, Kevin Y. Cho, Gary J. Patti, Brian N. Finck

## Abstract

Hepatic stellate cells (HSC) are non-parenchymal liver cells that produce extracellular matrix comprising fibrotic lesions in chronic liver diseases. Prior work demonstrated that mitochondrial pyruvate carrier (MPC) inhibitors suppress HSC activation and fibrosis in a mouse model of metabolic dysfunction-associated steatohepatitis (MASH). In the present study, pharmacologic or genetic inhibition of the MPC in HSC decreased expression of markers of activation *in vitro*. MPC knockdown also reduced the abundance of several intermediates of the TCA cycle, and diminished α-ketoglutarate played a key role in attenuating HSC activation by suppressing hypoxia inducible factor-1α signaling. On high fat diets, mice with HSC-specific MPC deletion exhibited reduced circulating transaminases, numbers of HSC, and hepatic expression of markers of HSC activation and inflammation compared to wild-type mice. These data suggest that MPC inhibition modulates HSC metabolism to attenuate activation and illuminate mechanisms by which MPC inhibitors could prove therapeutically beneficial for treating MASH.

## BACKGROUND and AIMS

The incidence of metabolic dysfunction-associated steatotic liver disease (MASLD; previously known as nonalcoholic fatty liver disease) and steatohepatitis (MASH) is growing rapidly. MASH is characterized by inflammation and fibrosis and dramatically increases the risk of developing cirrhosis, liver failure, and hepatocellular carcinoma [1–4]. Despite being a leading cause of liver-related morbidity and mortality, there are no licensed drug therapies for MASH and the molecular mechanisms driving the progression of MASLD to MASH are unclear [5]. The liver is composed of a variety of cell types including parenchymal (hepatocytes) and non-parenchymal cells (endothelial, immune, and stellate cells). While multiple cell lineages contribute to MASH progression, hepatic stellate cells (HSC) are the primary drivers of fibrosis in MASH and a variety of chronic liver diseases [6].

In their inactive (quiescent) state, HSC display an adipogenic-like transcriptional program and are one of the body’s major storage sites for vitamin A in the form of retinyl ester-rich lipid droplets [7]. In response to hepatic injury, HSC can proliferate, migrate to sites of injury, and differentiate into myofibroblasts that secrete components of the extracellular matrix that make up fibrotic lesions. HSC are activated in response to a variety of stimuli including factors released in response to cell injury and death, changes in the extracellular matrix, and signals from hepatocytes and immune cells including numerous cytokines and growth factors [8]. Although important for tissue repair, chronic activation of HSC leads to excess extracellular matrix protein (e.g. collagen I and collagen III) deposition in the liver, resulting in hepatic scarring (fibrosis) [9].

Importantly, activated HSC undergo dramatic changes in metabolism to meet the high demand for energy necessary for proliferation and ECM production [10]. Glucose uptake is enhanced and there are marked increases in rates of anaerobic glycolysis and mitochondrial pyruvate utilization upon HSC activation [11]. Activated HSC are also highly reliant on glutaminolysis as a fuel source [12]. Prior work has demonstrated that enhanced glucose uptake and glycolysis is required for fibroblasts to sustain TGFβ-induced collagen synthesis [13, 14] and interventions to suppress these metabolic changes have been shown to result in reduced HSC activation [11, 15]. Based on the importance of energy metabolism in HSC function, it seems plausible that these pathways may be targeted therapeutically to limit the development and progression of fibrosis.

Previous work from our lab has suggested that inhibitors of the mitochondrial pyruvate carrier (MPC) can suppress markers of HSC activation and reduce fibrosis in a mouse model of MASH [16]. The mitochondrial pyruvate carrier (MPC) plays a pivotal role in mitochondrial metabolism by transporting pyruvate, a product of glycolysis, into the mitochondrial matrix to enter the TCA cycle. The MPC is a heterodimeric complex composed of the MPC1 and MPC2 proteins [17, 18]. Both proteins are required for MPC stability, as the deletion of either protein destabilizes the complex [17–19]. Previous work has shown that MPC inhibition in multiple tissues provides metabolic benefits [19, 20] and reduces MASH progression in mice [16], but little is known about the direct effects of MPC inhibition on HSC metabolism and activation. To investigate this, we examined how disrupting the MPC affects stellate cell metabolism and activation *in vitro*, and generated mice with HSC-specific *Mpc2* deletion, to understand the effects *in vivo*. Here we show that pharmacologic or genetic inhibition of MPC attenuates the activation of hepatic stellate cells due to impaired mitochondrial αKG synthesis, which is required for HIF1α signaling pathway and collagen hydroxylation. Furthermore, we demonstrate that MPC deletion in HSC protects mice from HSC activation and inflammation in the context of a high fat, fructose, and cholesterol diet.

## RESULTS

### Inhibition of the MPC attenuates hepatic stellate cell activation in vitro

To test the role of MPC inhibition on HSC activation, we isolated HSC from the liver of wild-type mice and treated them with the MPC inhibitor (MPCi), 7ACC2 [21]. Compared to cells harvested one day after isolation, culturing isolated stellate cells for a period of seven days resulted in a substantial increase in several markers of HSC activation including multiple collagen isoforms, and non-collagenous enzymes that participate in ECM remodeling including *Spp1* and *Timp1* (Fig. 1a). However, treatment with 7ACC2, at doses previously shown to be specific towards the MPC [21], led to a significant reduction in several activation markers including *Col1a1*, *Col1a2*, *Spp1*, and *Timp1* (Fig. 1a).

**Fig. 1:**
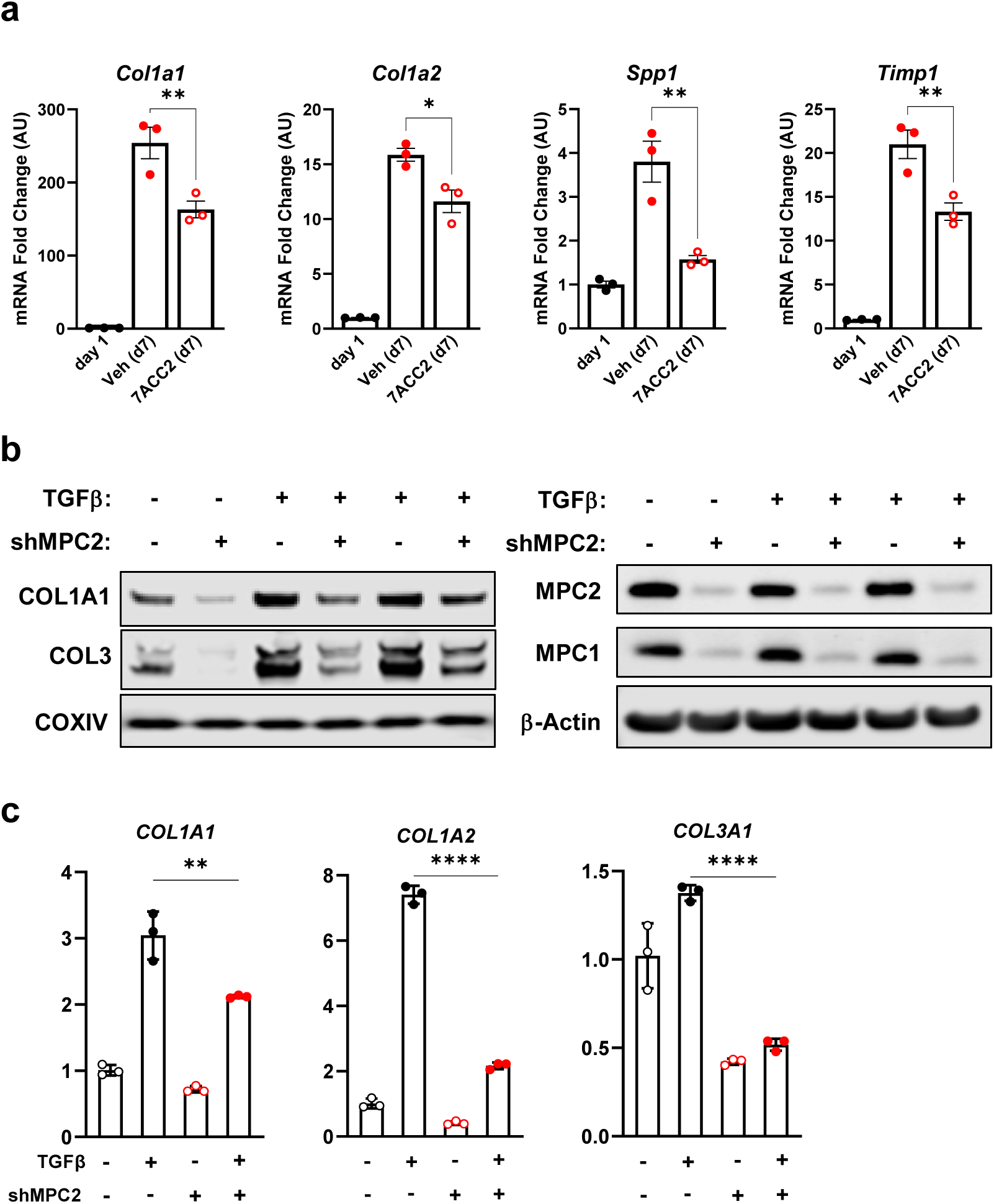
Genetic or pharmacological inhibition of the MPC reduces hepatic stellate cell activation in vitro. **a**, Hepatic stellate cells (HSC) were isolated from wild-type Mpc2^fl/fl^ mice and cultured for up to 7 days (d7) and treated with the addition of vehicle (DMSO; Veh) or 7ACC2 (1 µM). A portion were harvested after 1 day of culture (non-treated, d1) for quiescent HSC. Gene expression was measured by qRT-PCR and data are expressed as mean ± SEM, relative to d1 HSC. *p<0.05, **p<0.01. **b**, Human hepatic stellate cells (LX2) expressing shRNA against *MPC2*, reduced collagen protein abundance. TGFβ-1 (5 ng/mL) was added into culture medium to activate LX2 cells. **c**, Collagen isoforms gene expression were decreased in activated LX2 cells expressing shRNA against *MPC2*. Data are expressed as mean ± SEM, relative to control cells expressing non-targeting shRNA. *p < 0.05, **p < 0.01, ***p < 0.001, ****p < 0.0001.

We also evaluated the effects of suppressing MPC activity in a human HSC cell line by generating LX2 cells stably expressing shRNA against MPC2. As was observed with pharmacologic MPC inhibition, LX2 cells with MPC knockdown exhibited lower expression of collagen mRNA and protein compared to scrambled control cells after stimulation with TGFβ (Fig. 1b and 1c). Collectively, our results demonstrate that genetic or pharmacological inhibition of the MPC reduces HSC activation *in vitro*.

### Metabolic effects of MPC inhibition in hepatic stellate cells

To characterize LX2 metabolism and the effects of MPC inhibition on relevant metabolic pathways, we administered uniformly labeled ^13^C-glucose to control or MPC2 knockdown cells and then evaluated incorporation of ^13^C into various metabolites. MPC shRNA did not affect the pool size of glycolytic end products including pyruvate, lactate, and alanine (Fig. 2a). In contrast, MPC2 suppression markedly reduced the total abundance of several TCA cycle intermediates, including acetyl-CoA, citrate, aconitate, α-ketoglutarate (αKG), and succinate (Fig. 2b), which is consistent with the critical role of mitochondrial metabolism of glucose (as pyruvate) in generating these intermediates (Fig. 2c).

**Fig. 2:**
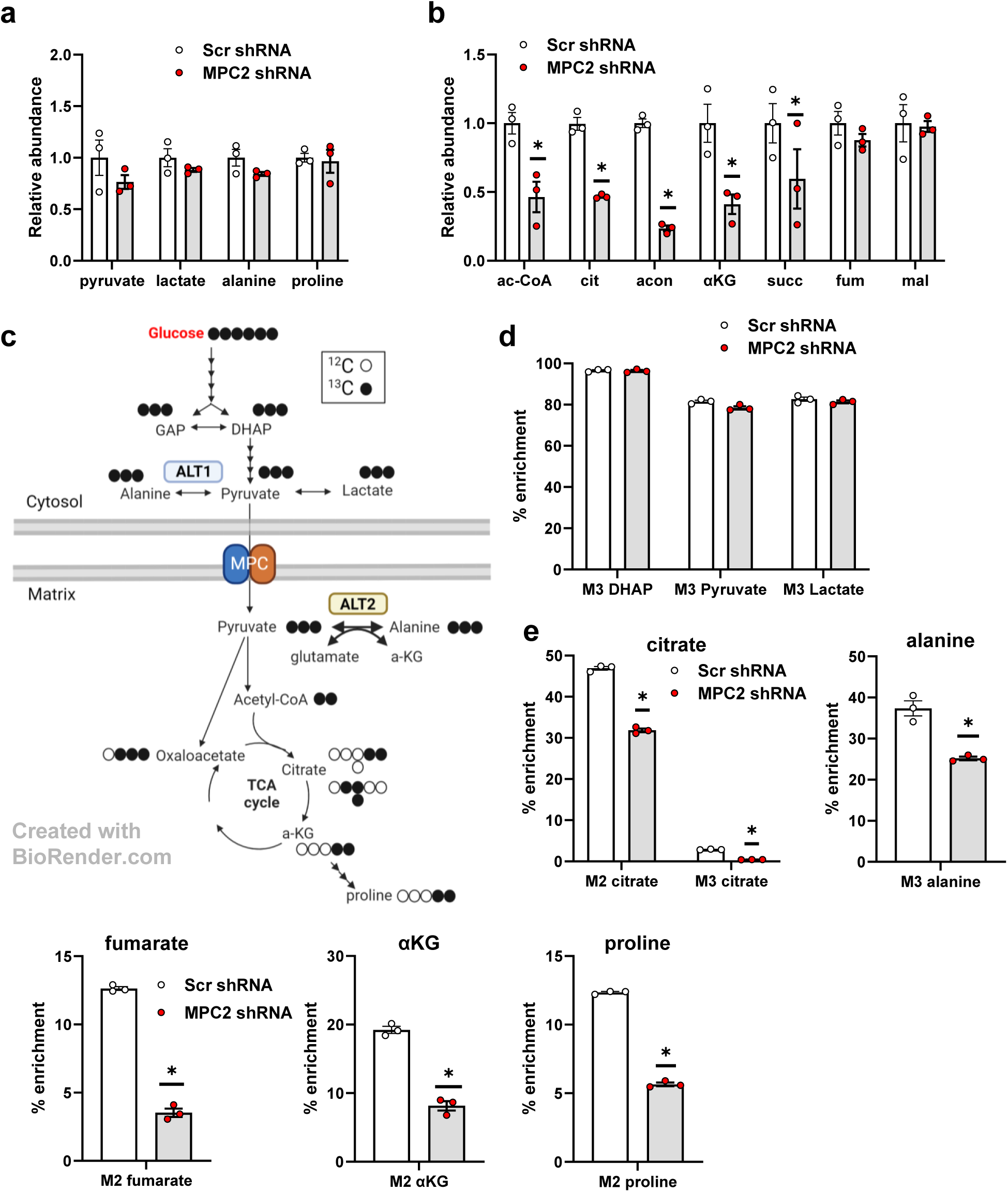
The MPC suppression markedly reduced several TCA cycle intermediates in LX2 cells treated with uniformly labeled ^13^C-glucose. **a**, The effect of MPC shRNA on relative abundance of glycolytic end products in LX2 cells treated with uniformly labeled ^13^C-glucose isotope. **b**, MPC suppression in LX2 cells reduced the relative abundance of several TCA cycle metabolites. **c**, Schematic of TCA cycle alterations measured by metabolomic analyses of LX2 cells stably expressing shRNA against *MPC2*. Red arrows, decreased; blue arrows, unchanged (comparing MPC2 shRNA to scrambled shRNA) **d**, Incorporation of ^13^C-glucose into glycolytic and TCA cycle intermediates in LX2 cells. TGFβ-1 (5 ng/mL) added into media to activate LX2 cells. GAP, glyceraldehyde 3-phosphate; DHAP, dihydroxyacetone phosphate; αKG, alpha-ketoglutarate; ALT2, alanine transaminase 2. Data are expressed as mean ± SEM, relative to TGFβ-1-free cells expressing non-targeting shRNA (n= 3 or 6). *p < 0.05.

Consistent with high rates of glycolysis in LX2 cells [22], ^13^C-glucose was abundantly incorporated into glycolytic intermediates including DHAP, pyruvate, and lactate, and this was not affected by MPC2 suppression (Fig. 2d). In contrast, MPC2 shRNA markedly reduced the incorporation of ^13^C into TCA cycle intermediates (citrate, fumarate, and αKG) and amino acids (alanine and proline) (Fig. 2e) that at least partially require mitochondrial import of pyruvate (Fig. 2c).

There were several interesting observations about the incorporation of glucose into TCA cycle intermediates and amino acids in LX2 cells. For example, by determining the number of ^13^C-labeled carbons from the abundance of M2-versus M3-labeled citrate, it is apparent that LX2 cells preferentially oxidize glucose-derived pyruvate rather than carboxylate it to oxaloacetate (Fig. 2e). Roughly 10% of the proline, a critical constituent of extracellular matrix as a component of mature collagen [23], in these cells was enriched from ^13^C-glucose (Fig. 2e). Suppression of MPC activity, markedly reduced incorporation of ^13^C-glucose into proline, consistent with a requirement for mitochondrial metabolism of pyruvate in this process. However, MPC2 suppression did not affect the total abundance of this amino acid (Fig. 2a), suggesting compensation by other metabolic pathways such as glutaminolysis.

We also observed that a high percentage (∼37%) of intracellular alanine was enriched in ^13^C from glucose (Fig. 2e). The conversion of pyruvate to alanine is mediated by the alanine transaminase (ALT) enzymes ALT1 and ALT2 encoded by genes called *GPT* and *GPT2*, respectively [24, 25]. ALT1 is cytosolic while ALT2 is found in the mitochondrial matrix (Fig. 2c). MPC2 knockdown significantly reduced the incorporation of ^13^C into alanine (Fig. 2e), suggesting that the mitochondrial ALT2 isoform contributes significantly to this metabolic fate. However, MPC deletion did not affect the total abundance of intracellular alanine (Fig. 2a).

### Role of alanine transaminase in pyruvate metabolism and HSC activation

To determine whether the reaction catalyzed by ALTs might be important for HSC activation, we treated LX2 cells with β-chloroalanine (BCLA), a transaminase inhibitor and then measured the expression of genes encoding collagens as a readout for HSC activation. We found that BCLA treatment attenuated LX2 cell activation in the context of TGFβ stimulation (Fig. 3a-b). Since BCLA inhibits both ALT enzymes, we also transfected LX2 cells with siRNA specific for *GPT2* (which encodes ALT2). *GPT2* siRNA resulted in suppression of *GPT2* mRNA and ALT2 protein abundance and suppressed the expression of *COL1A1* and a smooth muscle actin (ASM1) in LX2 cells (Fig. 3c). When MPC2 knockdown LX2 cells were transfected with *GPT2* siRNA, the combined knockdown of *MPC2* and *GPT2* had an additive effect on *COL1A1* mRNA.

**Fig. 3:**
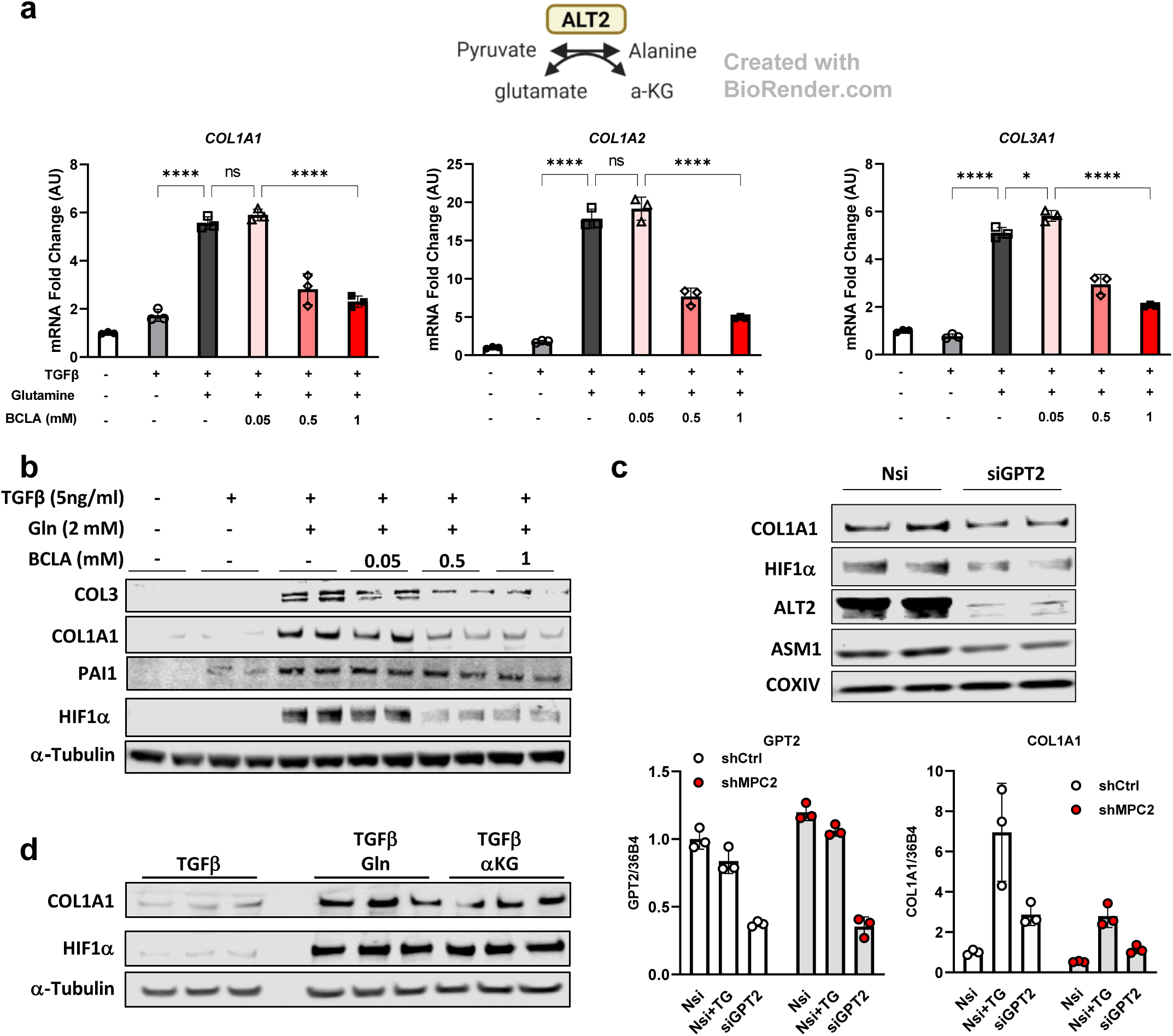
Alanine transaminase inhibition reduced HSC activation. **a**, Schematic for the reaction catalyzed by ALT2. *COL1A1*, *COL1A2*, and *COL3A1* gene expression in LX2 cells treated with/without TGFβ (5 ng/mL), glutamine (2 mM), and β-chloroalanine (BCLA; 0.05, 0.5, 1 mM). **b**, Western blot images for COL3, COL1A1, PAI1, HIF1α, and α-Tubulin in LX2 cells treated with BCLA. **c**, Protein abundance and gene expression of activation markers in either GPT2 knockdown LX2 or GPT2/MPC2 knockdown LX2 cells. For protein abundance, all groups were treated with 5 ng/mL TGFβ-1. **d**, Western blot images for COL1A1, HIF1α, and α-Tubulin in LX2 cells treated with or without glutamine (Gln, 2 mM) or dm-αKG (5 mM) compared with TGFβ-1 (5 ng/mL) treated cells.

To determine whether alanine availability might be sufficient to stimulate collagen gene expression, LX2 cells were kept in alanine/glutamine-free media or provided with increasing concentrations of alanine or 2 mM glutamine (positive control). Whereas glutamine was sufficient to activate *COL1A1, COL1A2,* and *COL3A1* gene expression, alanine was without effect (Supplemental Fig. 1a), suggesting that alanine does not play a critical role in the activation of LX2 cells.

The reaction catalyzed by ALT enzymes not only converts pyruvate to alanine, but also requires concomitant conversion of glutamate to αKG (Fig. 3a) [24, 25]. Metabolomic analyses demonstrated that αKG was depleted in cells with MPC knockdown (Fig. 2b). This is consistent with reduced synthesis through the ALT reaction and in the TCA cycle; both of which could be affected by loss of mitochondrial pyruvate import (Fig. 3a). We therefore tested the sufficiency of αKG to stimulate collagen expression by depriving cells of glutamine and then providing them with increasing amounts of a cell permeable αKG analog, dimethyl-αKG (dm-αKG). Interestingly, administration of dm-αKG to LX2 cells enhanced collagen protein expression similar to glutamine (Fig. 3d). These findings suggest that MPC knockdown alters the fate of glucose in LX2 cells and leads to depletion of αKG, which may be critical for HSC activation.

### MPC suppression leads to diminished activation of hypoxia inducible factor 1a

The hypoxia-inducible factor-1α (HIF1α) transcription factor may play important roles in connecting metabolic changes to HSC activation. HIF1α is a critical regulator of glycolytic enzyme expression [22, 26, 27], but also transcriptionally regulates several genes encoding fibrotic markers including *COL1A1* [28, 29], and is stimulated during the process of HSC activation [30]. Prior research has suggested that αKG might enhance the activation of HIF1α in lung fibroblasts [27]. To determine whether HIF1α was affected by αKG in our system, we measured HIF1α protein abundance in the presence or absence of dm-αKG or glutamine (positive control). The addition of either dm-αKG or glutamine markedly increased the abundance of HIF1α (Fig. 3d). Also consistent with αKG serving as the key trigger for HIF1α activation, treating cells with BCLA or siGPT2 suppressed the glutamine-induced increase in HIF1α abundance (Fig. 3b-c).

To determine whether HIF1α was required for the activation of collagen gene expression in response to these metabolic stimuli, we treated cells with several doses of the HIF1α inhibitor, CAY10585 [31]. The effects of glutamine or dm-αKG on COL1A1 protein abundance required HIF1α activity since addition of the HIF1α inhibitor CAY10585 suppressed this response to dm-αKG (Fig. 4a) or glutamine (Supplemental Fig. 1b).

**Fig. 4:**
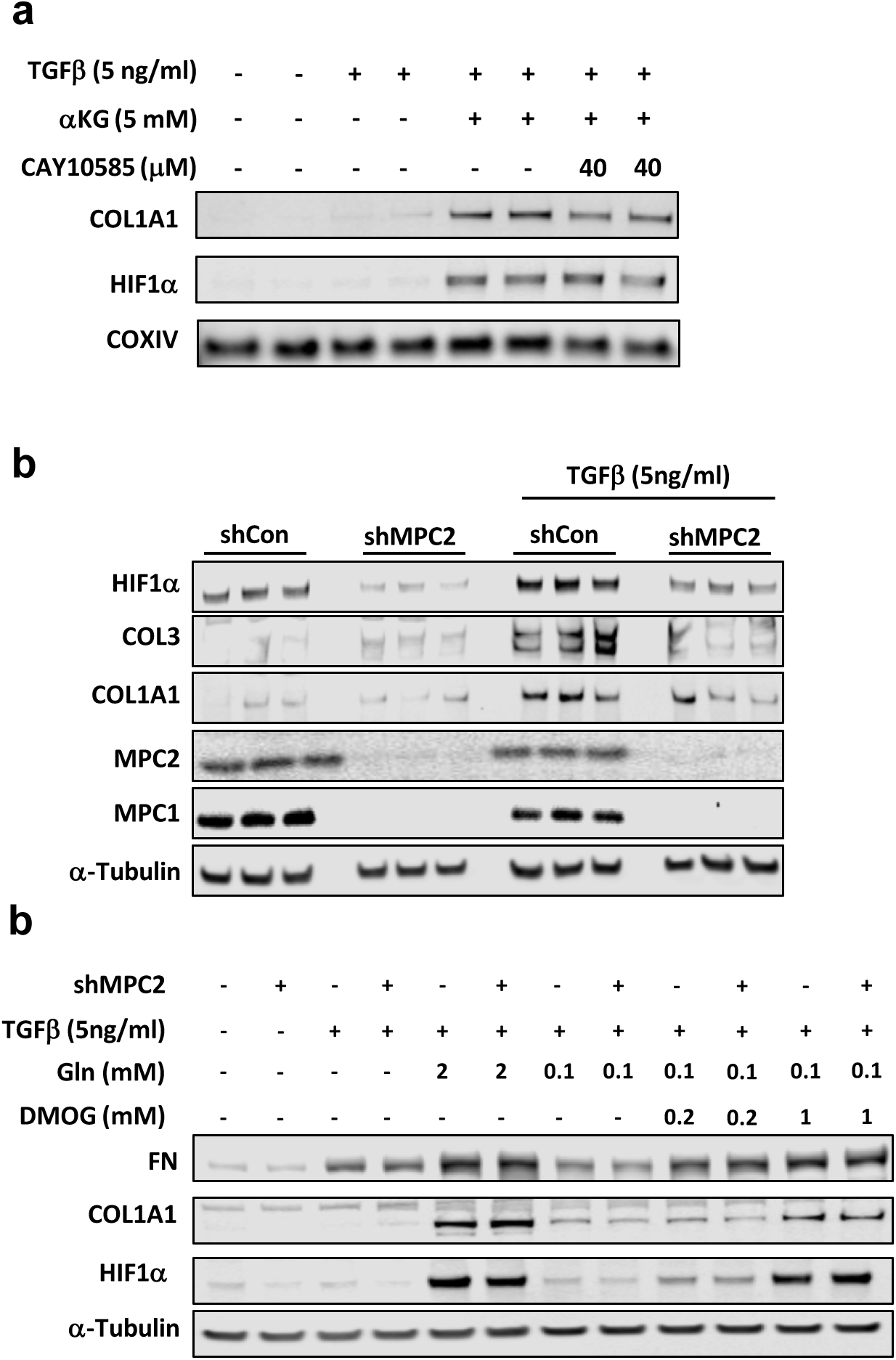
MPC suppression attenuated HSC activation through diminished activation of hypoxia inducible factor 1α. **a**, Western blot images for COL1A1, HIF1α, and COXIV in LX2 cells treated with dm-αKG (5 mM) and CAY10585. **b**, Western blot images for HIF1α, COL3, COL1A1, MPC2, MPC1, and α-Tubulin in LX2 cells expressing shRNA against *MPC2*. Cells treated with or without TGFβ-1 (5 ng/mL). **c**, Western blot images for FN, COL1A1, HIF1α, and α-Tubulin in LX2 cells expressing *MPC2* shRNA. Cells treated with TGFβ-1 (5 ng/mL), two doses of Gln (2 and 0.1 mM), and two doses of dimethyloxallyl glycine (DMOG; 0.2 and 1 mM;), a cell-permeable HIF1α stabilizer.

Next, we examined the effects of MPC inhibition on HIF1α abundance and found that shMPC2 expression reduced HIF1α abundance concordant with reduced collagen protein abundance (Fig. 4b). However, *HIF1A* mRNA was not affected by shMPC2 expression in LX2 cells (Supplemental Fig. 1c), consistent with a post-translational mechanism. Lastly, we treated LX-2 cells expressing shMPC2 or control shRNA with dimethyloxallyl glycine (DMOG) to stabilize HIF1α and found that DMOG overcame the effects of shMPC2 on HIF1α, COL1A1, and fibronectin (FN) protein abundance (Fig. 4c). Collectively, these findings suggest that suppression of MPC activity in LX2 cells attenuates activation by reducing αKG availability to affect HIF1α signaling.

### Lrat-Mpc2-/-mice are protected from MASH-inducing diet

To further investigate the role of the MPC in HSC, we generated mice with stellate cell-specific deletion of *Mpc2* by crossing *Mpc2^fl^*^/*fl*^ mice with mice expressing Cre recombinase under the lecithin-retinol acyltransferase (Lrat-Cre) promoter [6] to generate Lrat-Mpc2^-/-^ mice. First, we isolated and cultured HSC from both Lrat-Mpc2^-/-^ and wild-type littermate mice. As before, seven days in culture led to a considerable increase in several HSC activation markers in both groups, but the expression of these markers were substantially reduced in the Lrat-Mpc2^-/-^ HSC compared to WT cultures (Supplemental Fig. 2a). Next, we sought to examine how MPC inhibition in stellate cells affects liver injury and HSC activation in vivo. Lrat-Mpc2^-/-^ and wild-type littermate mice were placed on either a low-fat diet (LFD) or a diet high in fat, fructose, and cholesterol (HFC) for a period of 12 weeks. To exacerbate liver injury, we also treated HFC-fed mice with a single dose of carbon tetrachloride after four weeks on diet. At initiation of diet administration and after 12 weeks of diet, Lrat-Mpc2^-/-^ mice had an average body weight that was slightly less than wild-type mice, but both genotypes gained a similar amount during dietary feeding (Fig. 5a-b). Lrat-Mpc2^-/-^ mice on the HFC diet had a significant reduction in liver weight, fat mass, and lean mass, compared to wild-type; however, when normalized to total body weight, there was no difference in between groups (Fig. 5b-d, Supplemental Fig. 3a). Next, we assessed plasma transaminases, ALT and AST, as markers of liver injury and found that on the HFC diet Lrat-Mpc2^-/-^ mice had a significant reduction in both ALT and AST levels by ∼58% and ∼48%, respectively, compared wild-type mice (Fig. 5f). Histological examination of H&E-stained liver sections revealed a slight trend towards reduced macrosteatosis, inflammation, and NAFLD activity score, but these did not reach statistical significance (Fig. 5g, Supplemental Fig. 3b). However, quantification of desmin staining did show a significant decrease in the HSC staining in Lrat-Mpc2-/-mice relative to WT mice, but only on the HFC diet (Fig. 5g-h). There were no changes in plasma or intrahepatic lipid concentrations (Supplemental Fig. 3c-d). Lastly, we measured hepatic gene expression and found that Lrat-Mpc2^-/-^ mice had significantly decreased expression of several markers of HSC activation, including multiple collagen isoforms, *Acta2*, *Timp1*, and *Spp1* (Fig. 5i). Taken together, our results suggest that *Mpc2* deletion in hepatic stellate cells reduces the activation of these cells and protects from MASH development in mice fed a high-fat diet.

**Fig. 5:**
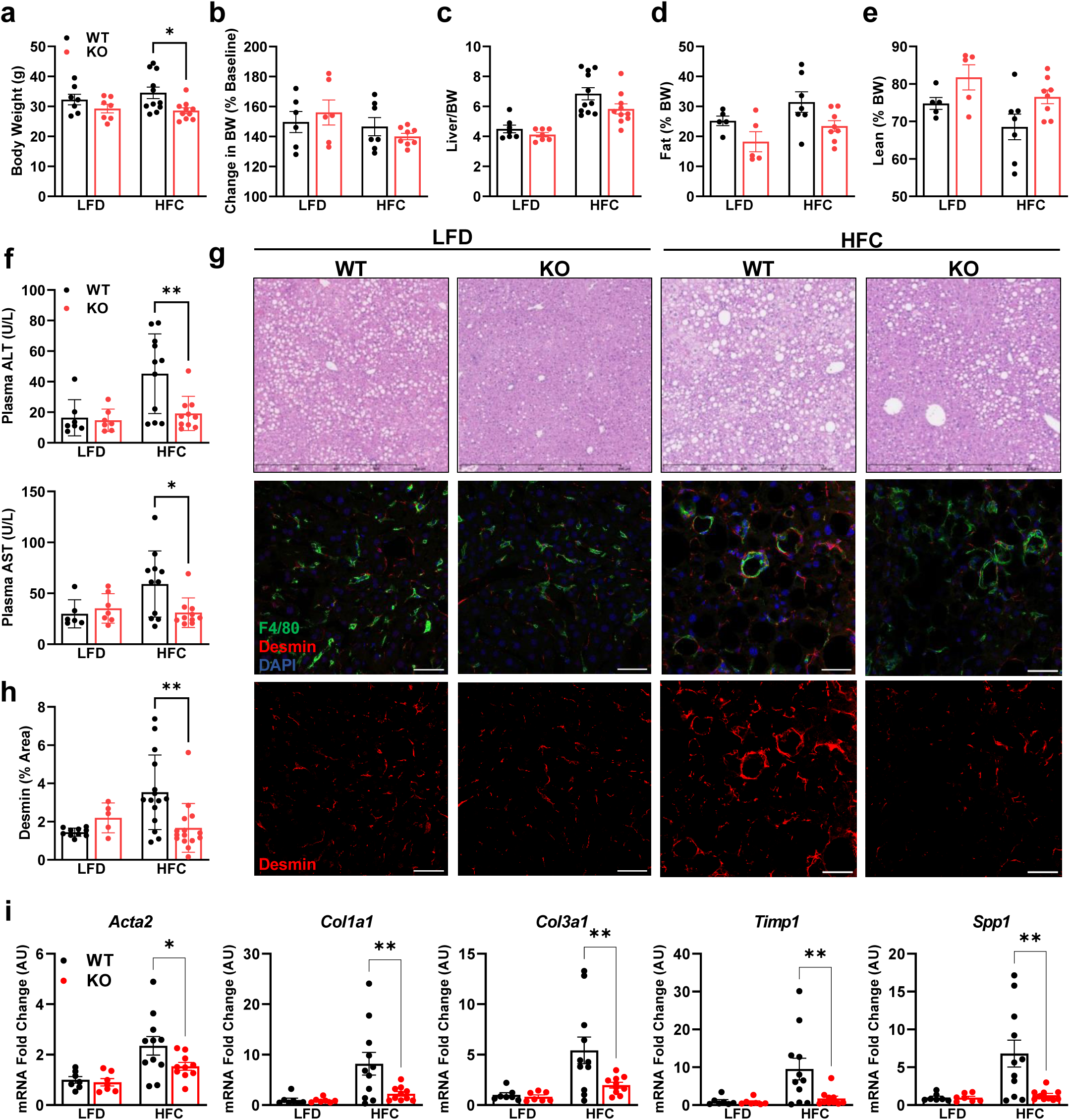
Lrat-Mpc2-/-mice are protected from MASH-inducing diet. At about 8 weeks of age, littermate wild-type (WT) and Lrat-Mpc2-/-(KO) mice were placed on either a low-fat diet (LFD) or a diet high in fat, fructose, and cholesterol (HFC) for a period of 12 weeks. **a,b**, Terminal body weight, expressed as total mass (**a**) and percent body weight gain from initiation of diet (**b**). **c**, Liver weight measured at sacrifice, expressed as percent body weight. **c**,**d**, Body composition determined by EchoMRI for both (**c**) fat mass and (**d**) lean mass, represented as percent body weight. **f**, Plasma levels of circulating transaminases ALT and AST collected at sacrifice. **g**, Representative liver sections with H&E staining and immunofluorescence staining for hepatic stellate cells (Desmin), macrophages (F4/80), and nuclei (DAPI). **h**, Quantification of desmin staining. **i**, Hepatic gene expression measured by RT-qPCR and expressed relative to WT LFD control group. All data expressed as mean ± SEM (n=7-11/group). *p<0.05, **p<0.01.

To gain a better understanding of the changes taking place in the hepatic transcriptome of Lrat-Mpc2^-/-^ and wild-type mice, we performed bulk RNA sequencing (RNAseq) analysis on livers from both genotypes on either the LFD or HFC diet. Analysis of differentially expressed genes (DEG) revealed Lrat-Mpc2^-/-^ mice had a decrease in the expression of 123 genes while 91 genes were increased compared to wild-type mice on the LFD (Fig. 6a). Comparison within the HFC groups revealed that Lrat-Mpc2^-/-^ livers exhibited reduced expression of 343 genes, while the expression of 133 genes was elevated, compared to wild-type mice (Fig. 6b). Gene set enrichment analysis a decrease in immune responses and apoptosis in Lrat-Mpc2^-/-^ relative to wild-type mice in both diet groups (Fig. 6c-d). Examination of perturbed KEGG signaling and metabolism pathways revealed a decrease in retinol metabolism and cell adhesion, while oxidative phosphorylation and protein processing were increased in Lrat-Mpc2^-/-^ mice on the LFD (Fig. 6e). On the HFC diet, Lrat-Mpc2^-/-^ mice again displayed a decrease in expression of immune signaling pathways while pathways associated with steroid hormone biosynthesis and amino acid metabolism were increased (Fig. 6f). Finally, pathway analyses also detected an increase in hypoxia signaling in WT mice with HFC diet and that was downregulated in Lrat-Mpc2^-/-^ mice compared to WT mice on the HFC diet (Fig. 6g-h). Indeed, we detected reduced expression of multiple HIF1α target genes in Lrat-Mpc2^-/-^ mice compared to WT mice on the HFC diet (Supplemental Fig. 4), which is consistent with our in vitro mechanistic studies described above. Collectively, these results indicate that stellate cell-specific deletion of *Mpc2* results in a shift in global hepatic transcriptome that is associated with a decreased immune response and possibly altered metabolism.

**Fig. 6:**
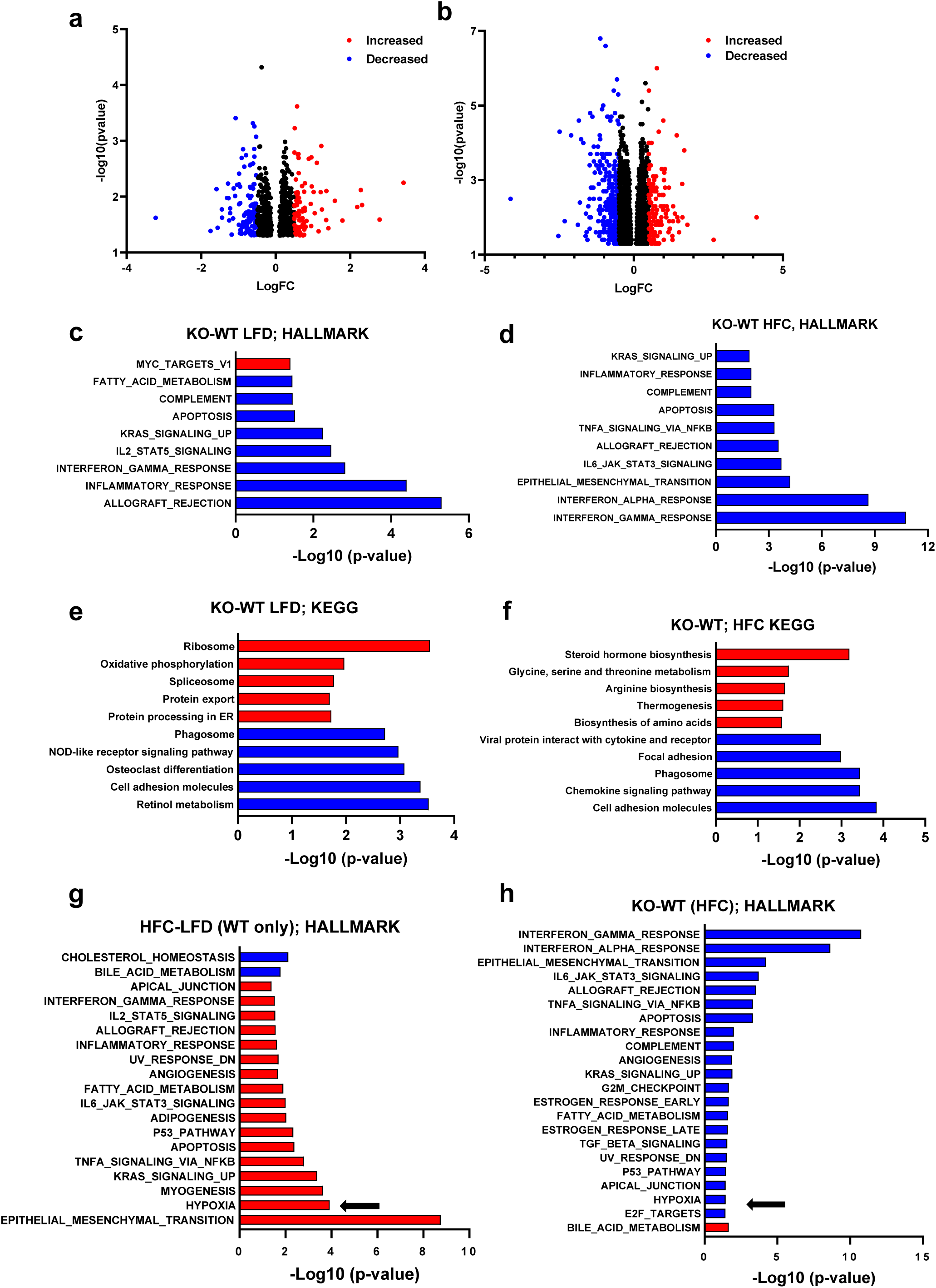
RNAseq reveals reduced immune responses and HIF1a signaling in Lrat-Mpc2-/-mice. RNA sequencing was performed on liver tissue from both wild-type (WT) and Lrat-Mpc2-/-(KO) mice and placed on either a LFD or HFC diet (n=5/group). **a**,**b**, Volcano plots of differentially expressed genes with p-value <0.05 comparing (**a**) KO versus WT mice on LFD and (**b**) KO versus WT mice on HFC diet. Differentially expressed genes with Log fold change (LogFC) less than –0.5, or greater than 0.5, were highlighted in either blue or red, respectively. Analysis of perturbations in Hallmark gene set collections when comparing (**c**) KO versus WT mice on LFD and (**d**) KO versus WT mice on HFC diet. Changes in KEGG signaling and metabolism pathways when comparing (**e**) KO versus WT mice on LFD and (**f**) KO versus WT mice on HFC diet. Analysis of perturbations in Hallmark gene set collections when comparing, **g**, HFC diet versus LFD in WT mice and, **h**, KO versus WT mice on HFC diet. The arrows highlight hypoxia signaling pathways.

### Lrat-Mpc2-/-mice are protected from MASH exacerbated by thermoneutral housing

To further confirm our findings and add scientific rigor, we used a second model to induce hepatic steatosis and injury by placing both Lrat-Mpc2^-/-^ and wild-type littermate mice in thermoneutral housing while feeding a western-type diet (42% kcal fat, 0.2% cholesterol) for a period of 20 weeks. Previous reports have demonstrated that housing at thermoneutrality, while feeding mice a western diet, exacerbates stellate cell activation and is associated with activation of inflammatory pathways that are like those found in human MASH development [32].

Like our previous experiment, Lrat-Mpc2^-/-^ mice had a modest but significant 15% decrease in body weight compared to wild-types, yet both groups gained a similar amount of weight over time (Fig. 7a). We again found that both ALT and AST levels were 57% and 43% lower, respectively, in Lrat-Mpc2^-/-^ mice compared to wild-type littermates (Fig. 7b). Total liver and fat mass were lower in Lrat-Mpc2^-/-^ mice compared to wild-type mice (Fig. 7c-d). As with our previous experiment, when liver and fat mass were normalized to body weight, there was no difference in liver or fat mass between genotypes (Fig. 7c-d). Lastly, Lrat-Mpc2^-/-^ mice had reduced hepatic expression of several markers of HSC activation including multiple collagen isoforms, *Timp1*, *Spp1*, and *Lgals3* (Fig. 7e). Collectively, our data demonstrate that stellate cell-specific disruption of the MPC blunts HSC activation and reduces markers of MASH development in mice.

**Fig. 7:**
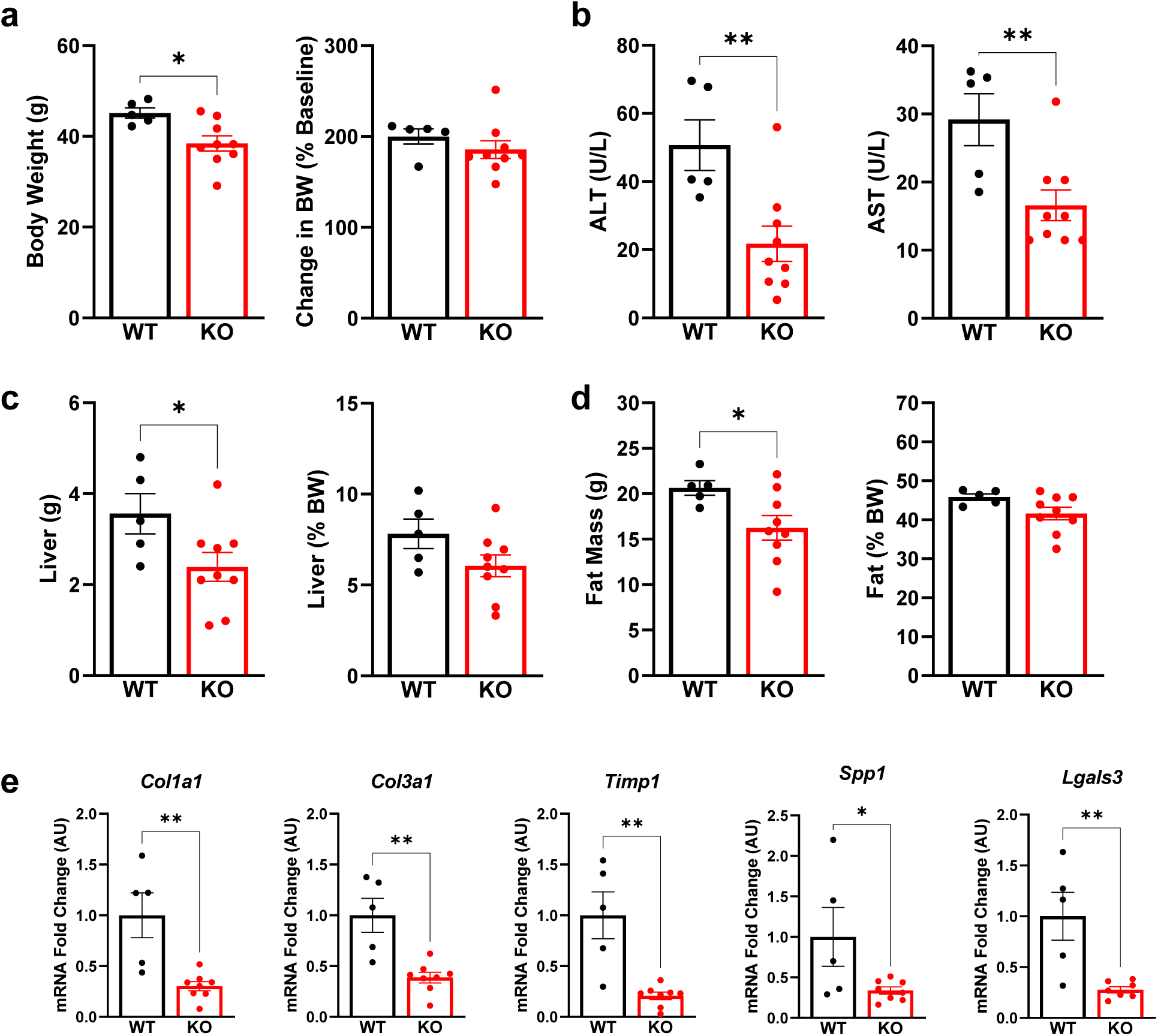
Lrat-Mpc2-/-mice are protected from MASH exacerbated by thermoneutral housing. At about 8 weeks of age, littermate wild-type (WT) and Lrat-Mpc2-/-(KO) mice were placed in thermoneutral housing (30°C) for 20 weeks. **a**, Terminal body weight, expressed as total mass and percent body weight gain from initiation of diet. **b**, Plasma levels of circulating transaminases ALT and AST collected at sacrifice. **c**, Liver weight measured at sacrifice, expressed as total mass and percent body weight. **d**, Analysis of fat mass determined by EchoMRI, represented as total mass and percent body weight. **e**, Hepatic gene expression measured by RT-qPCR and expressed relative to WT group. All data expressed as mean ± SEM (n=5-9/group). *p<0.05, **p<0.01.

## DISCUSSION

Activated HSC undergo complex changes in intermediary metabolism that allow them to proliferate, differentiate into myofibroblasts, and migrate to sites of injury. Prior work has demonstrated rates of glycolysis or glutaminolysis are markedly increased during the activation process and inhibiting flux through these metabolic pathways can attenuate HSC activation in response to activating stimuli [11, 15]. However, less is known about other pathways of intermediary metabolism and the therapeutic potential of targeting these pathways to attenuate HSC activation. In the current work, we sought to determine the effects of inhibiting the MPC, which connects anaerobic glycolysis with mitochondrial metabolism by transporting pyruvate, a major end-product of glycolysis, into the mitochondrial matrix where it can enter into the TCA cycle. Use of MPCi has been previously shown to reduce markers of HSC activation and fibrosis in a mouse model of MASH [16], but the direct effects on HSC activation has been little studied. We found that MPC pharmacological inhibition and genetic deletion of Mpc2, resulted in reduced activation of HSC *in vitro* and *in vivo.* We also determined that reduced αKG synthesis and abundance likely plays a mechanistic role in the inhibitory effects by suppressing HIF1α signaling. Importantly, we found that LratCre-Mpc2^-/-^ mice were also protected from liver injury and HSC activation in two high fat diet mouse models. Overall, these studies suggest that MPC inhibition can reduce HSC activation and suggest that inhibitors of this complex may have utility to prevent or reverse fibrosis in MASH and other chronic liver diseases.

Activated HSC undergo a dramatic shift in metabolism whereby there is an increase in glucose uptake and glycolysis, a phenomenon that is often found in cancer cells [10], and previous work has demonstrated that inhibition of anaerobic glycolysis reduces the activation of HSC [11]. However, very little is known about glucose oxidation (as pyruvate) in HSC. Herein, we show that ^13^C-labeled carbons in glucose become highly enriched in TCA cycle intermediates in activated HSC cells, suggesting that mitochondrial metabolism of pyruvate is robust. Indeed, inhibition of the MPC, which is required for mitochondrial pyruvate import to connect glycolysis and the TCA cycle, markedly reduced the pool size (relative abundance) and ^13^C-enrichments of several TCA cycle intermediates. One key metabolite reduced by MPC knockdown was αKG, which has been shown in other types of fibroblasts to play a key role in the activation process [27, 33–36]. αKG can be produced from a variety of metabolic processes including during oxidative metabolism of pyruvate in the TCA cycle, pyruvate transamination to alanine by transaminases, and through the process of glutaminolysis (Fig. 2c). Prior work has shown that addition of dm-αKG, a cell permeable form of αKG, was able to rescue HSC activation in the presence of the various glutaminolysis inhibitors [15]. In the present study, dm-αKG also reversed the effects of MPC knockdown (Fig. 4b) and inhibition of αKG synthesis by the ALT reaction suppressed COL1A1 expression even in the presence of glutamine (Fig. 3b).

The effects of αKG and MPC knockdown on HSC activation seem to be mediated, at least in part, via inhibition of HIF1α signaling. HIF1α is well known as a sensor of hypoxic conditions and enhances the expression of enzymes involved in anaerobic glycolysis as an adaptive response to limited oxygen availability [26]. However, HIF1α is also activated in response to other stimuli and also directly regulates the expression of a number of key genes involved in HSC activation [30]. HIF1α activity can be regulated at multiple levels including at the level of protein stability [37–39]. Under normoxic conditions, HIF1α protein is destabilized by prolyl hydroxylases (PHD), which are αKG-dependent dioxygenases [40, 41]. However, it has been shown that HIF1α abundance increased in human fibroblasts treated with dimethyl-αKG [42]. It might be expected that low αKG content would reduce PHD activity and therefore stabilization of HIF1α. The present finding that addition of αKG leads to greater HIF1α protein abundance is therefore somewhat counterintuitive. However, there are other examples where αKG depletion did not lead to HIF1α stabilization [43] and several of the present findings and other work conducted in fibroblasts [27, 42] are congruent with regards to αKG enhancing HIF1α abundance and fibroblast activation. This suggests that other mechanisms are at play and this should be examined in future studies.

Based on available data, we postulate that inhibition of the MPC limits HSC activation by reducing the ability of cells to use pyruvate as a metabolic substrate, which impacts signaling and biosynthetic processes. Although this work focused on the role of reduced αKG availability with MPC knockdown, other metabolic mechanisms may also be involved and should be assessed in the future. Indeed, MPC inhibition limited intramitochondrial pyruvate available for generation of acetyl-CoA, which may play a role in epigenetic regulation of gene expression. Alternatively, or in addition, limiting mitochondrial pyruvate metabolism may alter the metabolic fate of glutamine to impact the process of activation. Glutaminolysis is known to be critical for the activation process in HSC due to generation of αKG, involvement in proline and other amino acid biosynthesis for collagen production and modification, and as a TCA cycle substrate [12, 44]. It is possible that by reducing pyruvate utilization as an energy substrate, higher rates of glutamine use as an anaplerotic substrate for the TCA cycle limit its use for other purposes required for HSC activation.

Consistent with our in vitro findings, the presented in vivo studies demonstrated that mice with stellate cell-specific deletion of *Mpc2* were protected many effects of feeding diets that produce some features of MASH. Indeed, Lrat-*Mpc2*-/-mice were exhibited reduced circulating ALT and AST, reduced hepatic inflammation, as well as decreased expression of numerous markers of HSC activation. They also exhibited diminished numbers of HSC based on staining for desmin, but only in the context of the HFC diet. This could suggest that the ability to proliferate and activate in response to the HFC diet is impaired. Analysis of RNAseq data revealed a striking decrease in gene sets associated with both innate and adaptive immune responses, a key finding since immune cell infiltration is one of the early signs of disease development [45], but the exact mechanisms driving this are not clear. Finally, consistent with our in vitro mechanistic studies, bulk RNAseq analyses detected a signature for reduced HIF1α activation in the liver. While this approach cannot determine which cell type(s) exhibit reduced HIF1α activation, it is possible that this is reflecting diminished HIF1α signaling in HSC cells of the liver.

In summary, the current work provides evidence that inhibition of the MPC ameliorates HSC activation by altering intermediary metabolism to affect HIF1α signaling. Using Cre-LoxP mediated recombination, we generated novel stellate cell-specific *Mpc2* knockout mice, which were protected from hepatic injury and HSC activation in two separate dietary models. Analysis of hepatic RNAseq data demonstrated that HSC-specific *Mpc2* deletion resulted in a decreased in pathways associated with immune cell activation and extracellular matrix remodeling. Overall, these data highlight an alternative mechanism by which MPC inhibitors could be a novel therapeutic option for people afflicted with MASH.

## MATERIALS AND METHODS

### Hepatic stellate cell isolation

Primary hepatic stellate cells were isolated from mice as previously described [16]. Briefly, following pronase and collagenase perfusion of the liver, hepatocytes were removed from the cell suspension using a short centrifugation (50×*g* for 2 min at 4 °C). Then HSC were purified using a density gradient centrifugation using Optiprep (Sigma) and cells were seeded on to standard tissue culture dishes in DMEM media (Gibco) containing 10 % fetal bovine serum, and 1× Pen/Strep. After 24 hours, a subset of cells was harvested for use as a quiescent HSC control group. In the remaining cells, media was switched to media containing either vehicle (DMSO) or the MPCi 7ACC2 (1 µM; MedChemExpress; HY-D0713). Media was replenished every other day.

### LX2 experiments

The LX2 human stellate cell line (LX2 cells; Millipore Sigma, SCC064) was maintained in DMEM (Gibco, 11965-084) supplemented with 2% fetal bovine serum (FBS), sodium pyruvate (1 mM), and Penicillin-Streptomycin (100 U/mL). Unless otherwise mentioned, for cell culture experiments, LX2 cells were treated overnight with indicated treatments in Gln-free DMEM (Gibco, 11960-051) containing 10% FBS, 1 mM sodium pyruvate, and 100 U/mL Penicillin-Streptomycin. Reagents and inhibitors used in these studies included: TGFβ (PeproTech; AF-100-21C), CAY10585 (Cayman Chemical; 10012682), BCLA (Sigma-Aldrich; C9033), dm-αKG (Sigma-Aldrich; 349631), and DMOG (MedChemExpress; HY-15893).

### Lentivirus transfection

Lentiviral vectors expressing shRNAs against human MPC2, and lentiviral control non-targeting vector expressing scrambled shRNA were developed by FenicsBIO (MD, USA). The shRNA sequences used in this study were: Lentiviral human MPC2 shRNA-1 (CMV)(Cat. HSH-518021-1),5’-AAGATACTCACTTGTAATTATT-3’, Lentiviral human MPC2 shRNA-2 (CMV)(Cat. HSH-518021-2), 5’-GCCAGACCTGCAGAAAAACTTA-3’, and lentiviral control shRNA (CMV)(Cat. SH-CMV-C01). Following the induction of LX2 cells with lentiviral MPC2-targeting and non-targeting control vectors, puromycin (Sigma-Aldrich, P4512) was administered for stable selection of the cells expressing the shRNAs.

### Immunoblotting

RIPA lysis buffer (Cell Signaling Technology; 9806) with protease/phosphatase inhibitor cocktail (Cell Signaling Technology; 5872) was used to extract protein from liver or cultured cells as previously described [46]. For in vitro samples, cells were washed 1X with ice-cold PBS and lysis buffer was added in each well. Cells were scraped, briefly sonicated, and centrifuged for 10 minutes at 14,000 RPM in 4°C. For liver samples, approximately 30 mg of liver was homogenized using a TissueLyser and stainless steel beads. Protein was quantified using BCA assay (Thermo Scientific; 23227). Then, samples were mixed with 4X sample buffer (Invitrogen; NP0007) with β-mercaptoethanol (BioRad; 1610710). Proteins were heat denatured, and loaded on NuPAGE precast 4─12% Bis-Tris gels (Invitrogen; NP0322BOX; NP0329BOX; NP04122BOX) and run with MOPS (Invitrogen; NP0001) orMES (Invitrogen; NP0002) buffer and transferred to PVDF membrane (Sigma-Aldrich; IPFL00010). Antibodies used included: COL1A1 (Cell Signaling Technology; 72026), COL3 (ProteinTech; 22734-1-AP), FN (Abcam; ab2413), HIF1-α (Cell Signaling Technology; 36169), ALT1 (Abcam; ab202083), ALT2 (Sigma-Aldrich; SAB1409901), MPC1 (Cell Signaling Technology; D2L9I), MPC2 (Cell Signaling Technology; D4I7G), PAI1 (Cell Signaling Technology; 27535), α-TUBULIN (Sigma-Aldrich; T5168), β-ACTIN (Cell Signaling Technology; 3700), and COXIV (Cell Signaling Technology; 11967). To obtain western blot images, a Licor system was used along with the Image StudioLite software.

### Isotope-tracing experiments

LX2 cells expressing shMPC2 or scrambled shRNA control were incubated overnight in DMEM containing 25 mM U-[^13^C]-glucose, 10% fetal bovine serum (FBS), 2 mM Gln, 1 mM sodium pyruvate, and 1% penicillin/streptomycin and treated with 5 ng/mL TGFβ-1. Samples were collected and prepared for intracellular metabolite measurement as previously described [46]. Briefly, a portion of media were collected and stored at –80 °C for further processing. Cells were washed one time with pre-warmed PBS and one time with pre-warmed HPLC-grade water.

Then, quenched cells with ice cold HPLC-grade methanol, were scraped and collected into Eppendorf tubes followed by s SpeedVac drying for 2-3 hours. After being reconstituted in 1 mL of ice cold methanol:acetonitrile:water (2:2:1), all samples were vortexed, frozen in liquid nitrogen, and sonicated for 10 min at 25°C for three consecutive cycles. The samples were then centrifuged at 14,000×*g* at 4°C after being kept at 20°C for at least 1 hour. Supernatants were transferred to fresh tubes and SpeedVac dried for 2-5 h. Using a BCA kit from ThermoFisher, the protein content of cell pellets was assessed in order to standardize across samples.

Following the drying of the supernatant, 1 mL of water:acetonitrile (1:2) was added per 2.5 mg of cell protein, as measured in pellets recovered after extraction. Samples were subjected to two cycles of vortexing and 10 min of sonication at 25°C. Finally, samples were centrifuged at 14,000×*g* and 4°C for 10 min, the supernatant was transferred to LC vials and kept at –80 °C until MS analysis.

Ultra-high-performance LC (UHPLC)/MS was performed as previously described [47] with a ThermoScientific Vanquish Horizon UHPLC system interfaced with a ThermoScientific Orbitrap ID-X Tribrid Mass Spectrometer (Waltham, MA). Hydrophilic interaction liquid chromatography (HILIC) separation was accomplished by using a HILICON iHILIC-(P) Classic column (Tvistevagen, Umea, Sweden) with the following specifications: 100 × 2.1 mm, 5 µm. Mobile-phase solvents were composed of A ═ 20 mM ammonium bicarbonate, 0.1% ammonium hydroxide and 2.5 µM medronic acid in water:acetonitrile (95:5) and B ═ 2.5 µM medronic acid in acetonitrile:water (95:5). The column compartment was maintained at 45 °C for all experiments. The following linear gradient was applied at a flow rate of 250 µL min^─1^: 0─1 min: 90% B, 1─12 min: 90-35% B, 12─12.5 min: 35-25% B, 12.5─14.5 min: 25% B. The column was re-equilibrated with 20 column volumes of 90% B. The injection volume was 2 µL for all experiments. Data were collected with the following settings: spray voltage, ─3.0 kV; sheath gas, 35; auxiliary gas, 10; sweep gas, 1; ion transfer tube temperature, 250 °C; vaporizer temperature, 300 °C; mass range, 67─1500 Da, resolution, 120,000 (MS1), 30,000 (MS/ MS); maximum injection time, 100 ms; isolation window, 1.6 Da. LC/ MS data were processed and analyzed with the open-source Skyline software [48].

### RNA isolation and quantitative real-time PCR

RNA was isolated as previously described [49]. For in vitro studies, RNA was harvested from tissue culture dishes using TRIzol Reagent (Ambion) and Purelink™ RNA Mini Kit (Invitrogen™). For liver tissue, ∼30 mg of frozen liver tissue was homogenized in TRIzol Reagent (Ambion) using the TissueLyser II (Qiagen) followed by isolation using the Purelink™ RNA Mini Kit (Invitrogen™). RNA was reverse transcribed into complimentary DNA (cDNA) using High-Capacity reverse transcriptase (Life Technologies).

Quantitative real-time polymerase chain reaction was performed using Power SYBR Green (Thermo Fisher Scientific) and an optical 384-Well Reaction Plate (Applied Biosystems) using ViiA 7 Real-Time PCR System (Applied Biosystems). Relative gene expression was determined by the ΔΔCt method *36b4* as a reference gene. The primer sequences applied for gene expression are available upon request.

### Animal Studies

All animal experiments were approved by the Institutional Animal Care and Use Committee of Washington University in St. Louis (animal protocol number 20-0004). For all animal experiments, male mice in the C57BL/6J background were used. To generate HSC-specific deletion of Mpc2, we crossed *Mpc2*^fl/fl^ mice with lecithin retinol acyltransferase-Cre (Lrat-Cre) mice as previously described [6]. Littermates that did not express *Cre* were used as controls in all studies. For high fat diet studies, 8-week-old male mice were fed with a low-fat diet (Research diets D09100304) or a diet high in fat, fructose, and cholesterol (Research diets D09100310) for a period of 12 weeks. Four weeks after the initiation of experimental diets (*i.e*., 12 weeks of age), mice received a single intraperitoneal injection of carbon tetrachloride (CCL4) dissolved in corn oil (0.5 µL CCL_4_ per gram body weight). Finally, after 12 weeks on diet, animals were fasted for 5 h, euthanized by using CO_2_ asphyxiation, and tissue samples were collected for further analyses. Blood samples were collected through inferior vena cava into EDTA-containing tubes.

For thermoneutral studies, mice were switched from a standard chow diet to a western-type diet (42% kcal fat, 0.2% cholesterol; TD88137). At the same time, mice were placed in thermoneutral housing (30 °C). Mice were maintained on dietary feeding and thermoneutral housing for 20 weeks then sacrificed for tissue and blood collection following a 5 h fast.

### Body composition

Total body composition of fat and lean mass was determined by EchoMRI-100H (EchoMRI LLC).

### Plasma analyses

Plasma levels of alanine transaminase (ALT) and aspartate transaminase (AST) were determined using kinetic absorbance assays (Teco Diagnostics) as described previously [16]. Analysis of plasma triglycerides (Thermo Fisher Scientific), total cholesterol (Thermo Fisher Scientific), and non-esterified fatty acids (FUJIFILM Wako) were determined using colorimetric based assay as previously reported [50, 51].

### Hepatic triglyceride and NEFA analyses

Frozen liver samples were weighed (50-100 mg) and homogenized in ice-cold PBS (Gibco, 14190-051). Then, hepatic lipids were solubilized in 1% sodium deoxycholate and using the commercially available kits, hepatic triglyceride (Thermo Scientific, TR22421) and NEFA (Fuji Film) were quantified according to the manufacturer’s directions.

### Histological Analyses

Liver tissue was fixed in 10% neutral buffered formalin for 24 hours then paraffin embedded, then sections were cut and stained with hematoxylin and eosin (H&E). H&E sections were assessed for steatosis score, lobular inflammation, and NAFLD total score, by a board-certified pathologist (M.H.) who was blinded from treatment groups, as previously reported [16, 51].

### Immunofluorescence and quantification

Livers were fixed in 10% formalin for 24h at 4°C, then washed and incubated in 30% sucrose for another 24h at 4°C. The specimens were then embedded in OCT compound, cut into 8-µm thick sections by a cryostat (Leica Biosystems; Wetzlar, Germany), and stored at –80°C. For staining, sections were air-dried for 12min, rehydrated in PBS for 5min, and then blocked with freshly prepared blocking buffer (PBS +1% BSA w/v +0.3% Triton X-100 v/v) for 1h at room temperature. After blocking, 50uL of primary antibody cocktail prepared in blocking buffer was deposited on the section for overnight incubation at 4°C in a humidified chamber. The antibody solution was then aspirated, and the sections were washed with PBS, 3 times, 5min each. The secondary antibody cocktail was then similarly deposited on the section and incubated for 1h at room temperature in the dark.

Sections were then washed 3 times with PBS, 5min each, and nuclei were stained with fresh Hoechst dye (1:25,000, in PBS) for 5 min in the dark at room temperature. The sections were then mounted with prolong gold antifade mounting reagent and a #1 coverslip. Antibodies used were: F4/80 (eBioscience 13-4801-85; 1:200), Desmin (Abcam Ab15200; 1:100), anti-Rat IgG AF488 (Invitrogen A21208; 1:500), and anti-Rabbit IgG (Jackson ImmunoResearch 711-585-152; 1:200).

Confocal images were acquired using an LSM 900 laser scanning confocal microscope (ZEISS; Jena Germany) with a 20× 0.8 N.A. objective at ambient temperature. All the images were acquired in the same period using the same laser intensity and gain. The brightness of individual channels was optimized uniformly in Fiji (ImageJ, National Institutes of Health, Bethesda, MD, USA) by adjusting minimum and maximum displayed values. Quantification on the desmin-positive area was performed blindly using manual thresholding. Five randomly chosen fields (excluding large vessels) per sample were acquired for quantification.

### RNA sequencing

RNA was isolated from liver tissue as described above and bulk RNA sequencing was performed at the Genomic Technologies and Access Center at the McDonald Genomic Institute of Washington University School of Medicine in St. Louis. An Agilent Bioanalyzer was used to assess RNA integrity and ribosomal RNA depletion was performed with RiboErase (HMR). Samples were prepared according to library kit manufacturer’s protocol, then indexed, pooled, and sequenced on a NovaSeq S4 2×150, targeting ∼30 million reads per sample. Detailed methods of RNA sequencing analysis are previously reported [49].

### Statistical Analyses

GraphPad Prism Software, version 9.5.0 for windows, was used to generate figures. All data and are presented as the mean ±SEM. Statistical significance was calculated using an unpaired Student’s t-test, two-way analysis of variance (ANOVA), or one-way ANOVA with Tukey’s multiple comparisons test, with a statistically significant difference defined as p<0.05.

## ACKNOWLEDGMENTS

This work was funded by NIH grant R01 DK104735 (to B.N.F.). The Core services of the Diabetes Research Center (P30 DK020579), Digestive Diseases Research Cores Center (P30 DK052574), and the Nutrition Obesity Research Center (P30 DK56341) at the Washington University School of Medicine also supported this work. D.F was supported by a training grant (T32 DK007120) and NIH K01 award DK137050. J.D.S. is supported by R01 DK131188. Some metabolic analyses were supported by NIH grant R35 ES028365 (G.J.P.). We would like to thank Dr. Robert Schwabe for providing Lrat-Cre transgenic mice and Dr. Scott Friedman for the gift of LX2 cells.

## FIGURE LEGENDS

**Supplemental Fig. 1:**
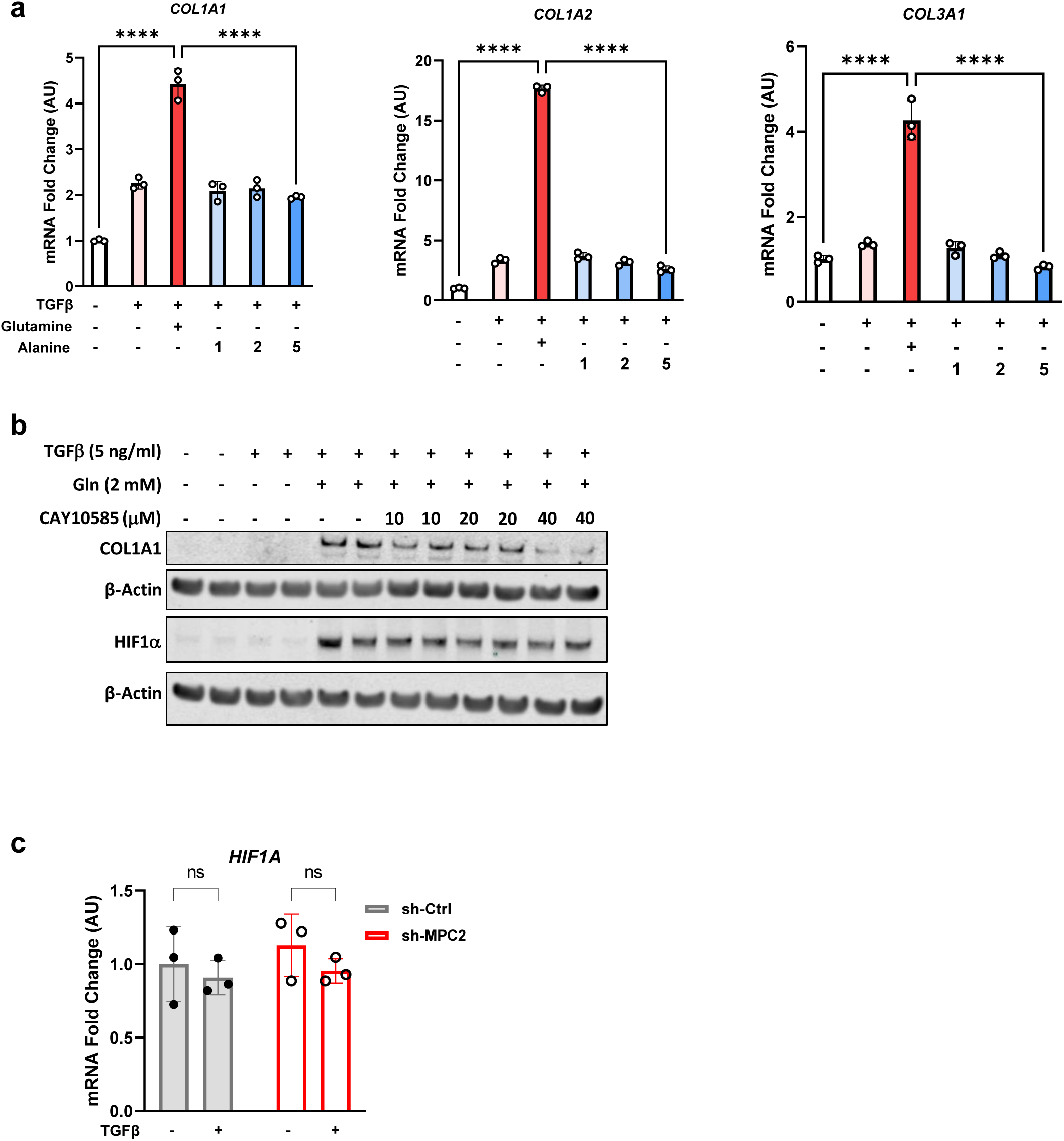
Alanine is not critical for the activation of hepatic stellate cells. **a**, LX2 cells activated by TGFβ-1 (5 ng/mL) and treated with or glutamine (2 mM) or three doses of alanine (1, 2, or 5 mM). *COL1A1*, *COL1A2*, and *COL3A1* gene expression measured by RT-qPCR and expressed as mean ± SEM, relative to TGFβ-1-free cells. **b,** Western blot images for COL1A1, HIF1α, and β-Actin in LX2 cells treated with Gln and HIF1α inhibitor, CAY10585. **c**, The mRNA abundance of *HIF1A* in TGFβ-1-activated LX2 cells expressing scramble or MPC2 shRNA and data expressed as mean ± SEM, relative to TGFβ-1-free scramble-shRNA cells. ns, non-significant, *p < 0.05, **p < 0.01, ***p < 0.001, ****p < 0.0001.

**Supplemental Fig. 2:**
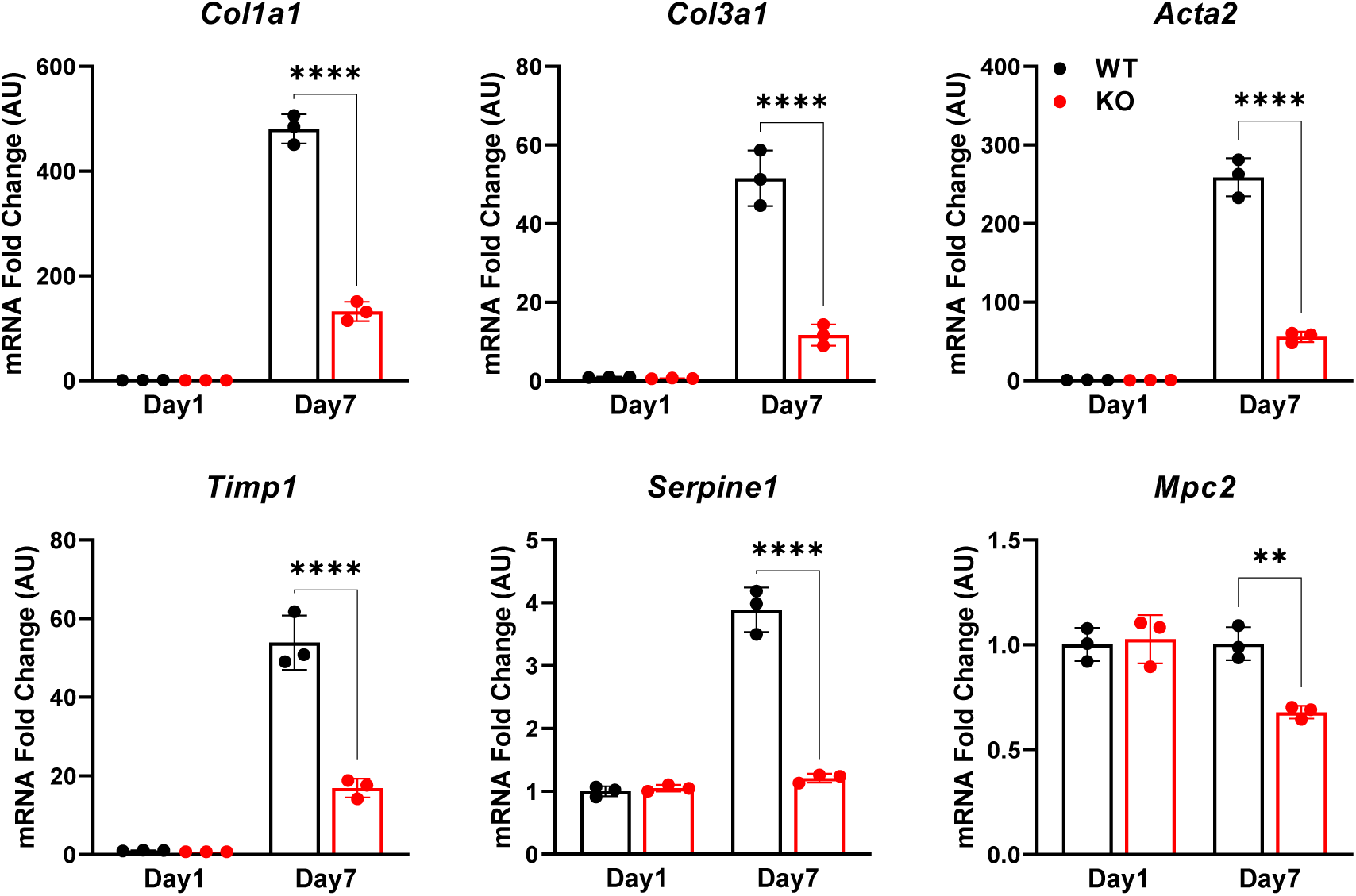
Stellate cell-specific deletion of *Mpc2* blunts HSC activation in vitro. *Col1a1*, *Col3a1*, *Acta2*, *Timp1*, *Serpine1*, and *Mpc2* gene expression of isolated hepatic stellate cells (HSC) from wild-type Mpc2^fl/fl^ mice and MPC2-/-littermates that cultured for up to 7 days (Day7). A portion were harvested after 1 day of culture (Day1) for quiescent HSC. Gene expression was measured by RT-qPCR and data are expressed as mean ± SEM, relative to day1 HSC. **p < 0.01, ****p < 0.0001.

**Supplemental Fig. 3:**
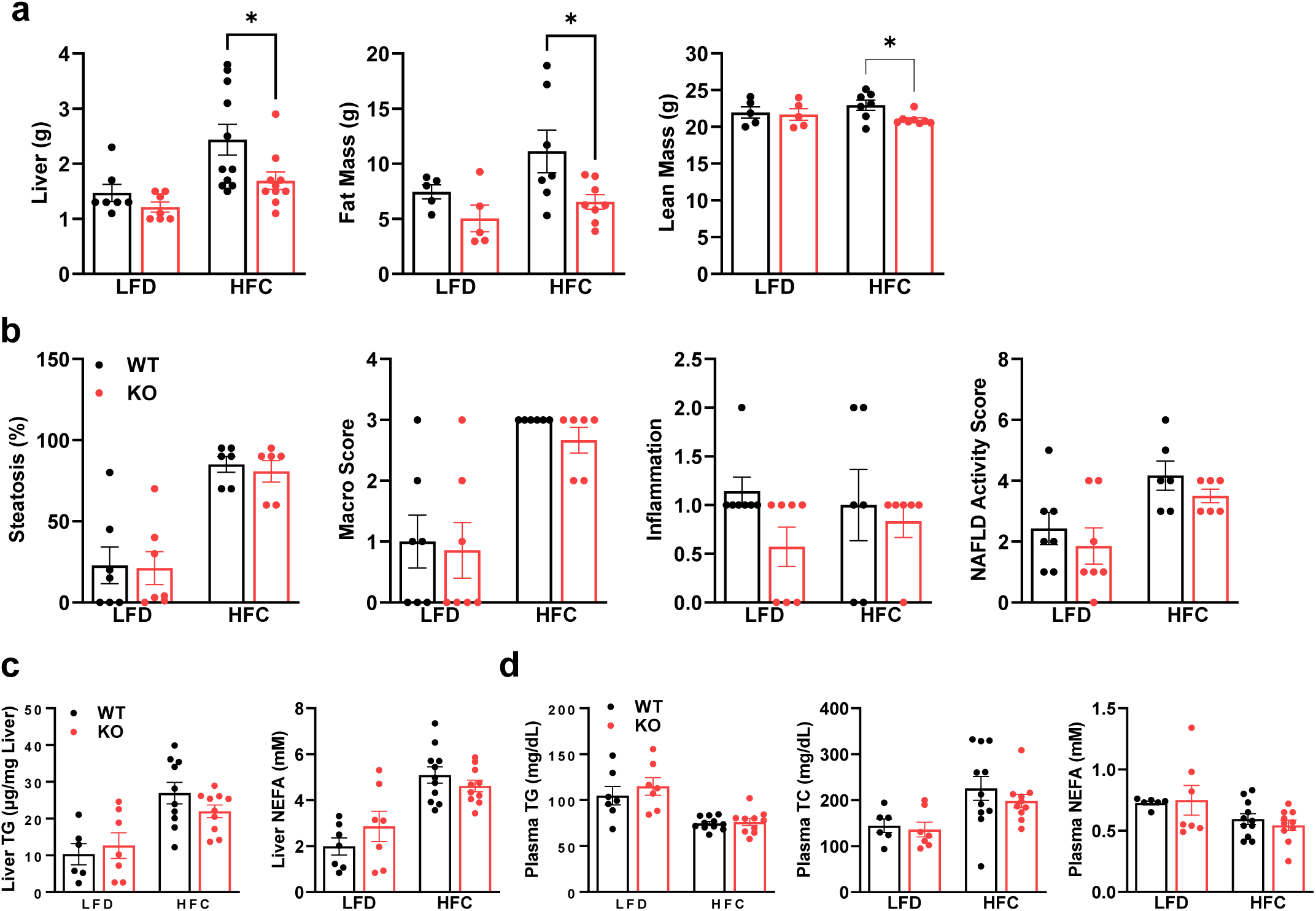
Plasma lipids were not affected in Lrat-MPC2-/-mice fed a MASH-inducing diet. At about 8 weeks of age, littermate wild-type (WT) and Lrat-MPC2-/-(KO) mice were placed on either a low-fat diet (LFD) or a diet high in fat, fructose, and cholesterol (HFC) for a period of 12 weeks. **a**, Liver weight, measured at sacrifice, and body composition, determined by EchoMRI, expressed as mean ± SEM (n=7-11/group). **b**, Histological scoring of H&E-stained liver sections assessing steatosis, macrosteatosis, lobular inflammation, and NAFLD activity score expressed as mean ± SEM (n=6-7/group). **c,d**, Analysis of triglycerides (TG), total cholesterol (TC), and non-esterified fatty acids (NEFA) from liver (**c**) and plasma (**d**), expressed as mean ± SEM (n=7-11/group).

**Supplemental Fig. 4:**
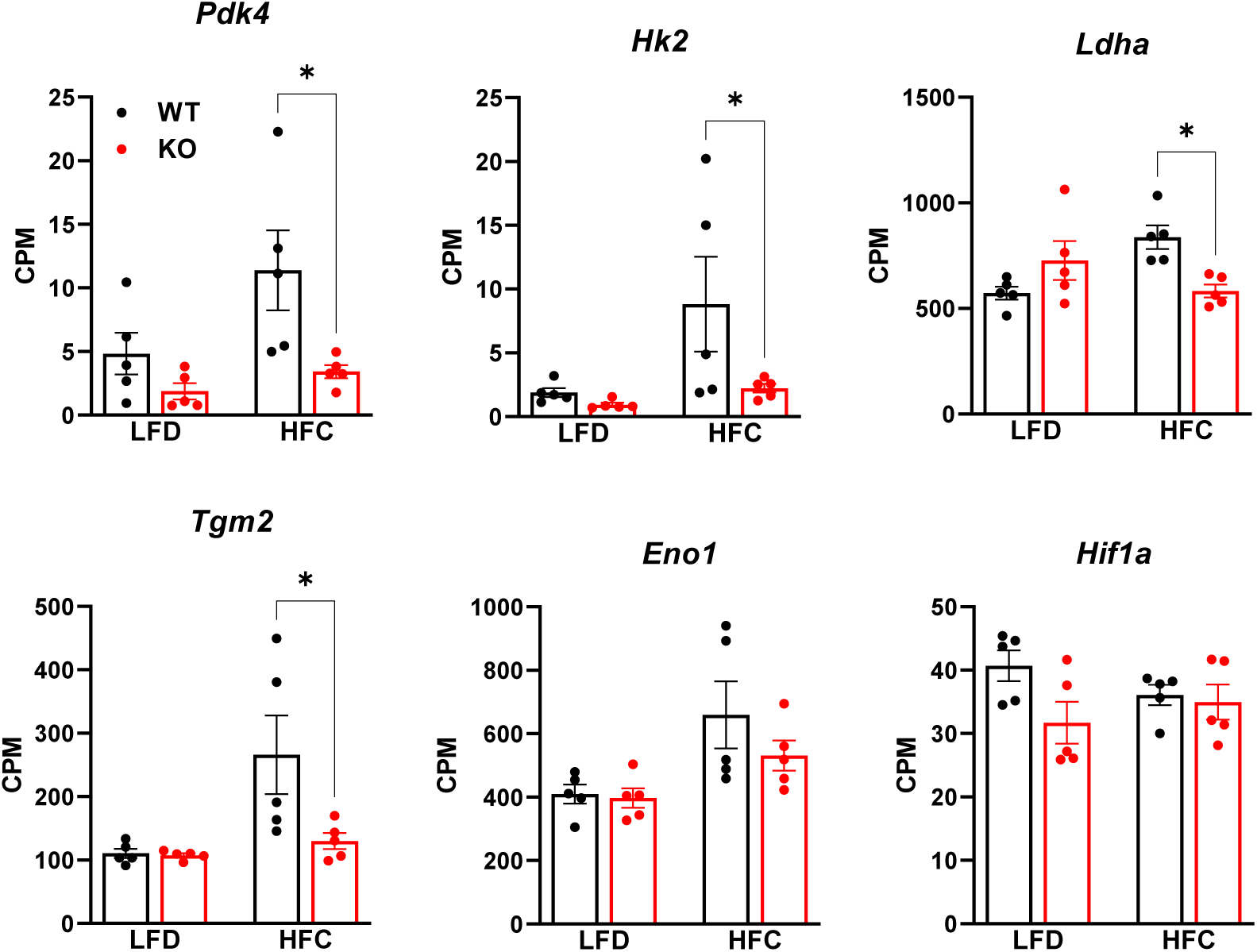
Diminished expression of HIF1α target genes in Lrat-Mpc2-/-mice on HFC diet. RNA sequencing was performed on liver tissue from both wild-type (WT) and Lrat-Mpc2-/-(KO) mice and placed on either a LFD or HFC diet (n=5/group). Selected HIF1α target genes from RNAseq data expressed as counts per million (CPM) and represented as mean ± SEM (n=5/group).

